# Near Infrared Diffuse *In Vivo* Flow Cytometry

**DOI:** 10.1101/2022.05.11.491330

**Authors:** Joshua Pace, Fernando Ivich, Eric Marple, Mark Niedre

## Abstract

**Significance:** Diffuse in vivo Flow Cytometry (DiFC) is an emerging technique for enumerating rare fluorescently labeled circulating cells non-invasively in the bloodstream. Thus far we have reported red and blue-green versions of DiFC. Use of near-infrared (NIR) fluorescent light would in principle allow use of DiFC in deeper tissues and would be compatible with emerging NIR fluorescence molecular contrast agents.

**Aim:** In this work, we describe the design of a NIR-DiFC instrument and demonstrate its use in optical flow phantoms *in vitro* and in mice *in vivo*.

**Approach:** We developed an improved optical fiber probe design for efficient collection of fluorescence from individual circulating cells, and efficient rejection of instrument autofluorescence. We built a NIR-DiFC instrument. We tested this with NIR fluorescent microspheres and cell lines labeled with OTL38 fluorescence contrast agent in a flow phantom model. We also tested NIR-DiFC in nude mice injected intravenously with OTL38-labeled L1210A cells.

**Results:** NIR-DiFC allowed detection of CTCs in flow phantoms with mean signal to noise ratios (SNRs) of 19 to 32 dB. In mice, fluorescently-labeled CTCs were detectable with mean SNR of 26 dB. NIR-DiFC also exhibited orders significantly lower autofluorescence and false-alarm rates than blue-green DiFC.

**Conclusions:** NIR-DiFC allows use of emerging NIR contrast agents. This work could pave the way for future use of NIR-DiFC in humans.

## 1 Introduction

In hematogenous cancer metastasis, circulating tumor cells (CTCs) intravasate from the primary tumor into the peripheral blood. A small fraction of these travel to distant organs via the circulatory system and form secondary tumors (1-6). As such, the ability to detect, count, and characterize CTCs and CTC clusters is of potentially great diagnostic value in clinical management of cancer and in the basic study of cancer biology (7-9). Fewer than 1 CTC per mL of peripheral blood may be indicative of metastatic progression (10-15). “Liquid biopsy” is the current gold standard for the study of CTCs, which involves drawing and analyzing small volume (∼mL) peripheral blood samples using ex-vivo assays such as *CellSearch* (16, 17). While powerful, liquid biopsy is poorly suited to measuring dynamic changes in CTC numbers over time (18, 19). In addition, because volume of peripheral blood is factionally very small compared to the total blood volume (<1%) these may provide a poor overall and statistical picture of CTC numbers (18-21).

These limitations have motivated development of optical methods to enumerate CTCs directly in circulation in vivo (22-27). Our team developed a pre-clinical laser-induced fluorescence technique called ‘Diffuse *in viv*o Flow Cytometry’ (DiFC) (28, 29). DiFC uses specially designed optical fiber probes and highly scattered light to sample blood flowing in large vessels in the tail or the hindleg of a mouse. By measuring the weak signal from emitted cells, DiFC allows non-invasive sampling of about 100 µL of blood per minute. We previously used DiFC to study CTC dissemination in preclinical mouse models including a subcutaneous Lewis lung carcinoma (LLC) and multiple myeloma (MM) disseminated xenograft model (30, 31). Because DiFC is non-invasive, the entire peripheral blood volume of a mouse can be sampled continuously and repeatedly over time.

Thus far, our DiFC instruments were designed to work with either blue-green (e.g. GFP) or red (e.g. Cy5) fluorophores that are commonly used in pre-clinical research. One long-term goal of our research is potential human translation of DiFC technology (32). Because of the fundamental nature of light transport in biological tissue, visible light is well suited to tissue depths of about 1 mm or less but is not well suited for use in larger limbs as would be required in humans.

In this paper, we describe the design and validation of a new, near-infrared (NIR)-DiFC instrument. NIR light is known to experience lower attenuation (scattering and absorption) in biological tissue than visible light, and therefore is expected to be better suited to larger anatomies (33). We recently showed that NIR light would enable detection of well-labeled CTCs in large blood vessels 2-4 mm below the surface, which would be suitable for measurement of cells in the human wrist or forearm (34, 35). Hence, development of NIR-DiFC is a logical first step in potential human translation of the technique.

Related to this, any use of DiFC in humans would also require the use of an administered molecular fluorescence contrast agent that would allow specific and bright labelling of CTCs. We and others have already showed that this is feasible using small-molecule or antibody targeted fluorescent molecular probes that are specific to cancer-associated cell surface markers (36-38). There are a large and growing number of molecular contrast agents under development for fluorescence guided surgery with NIR dyes including indocyanine green (ICG), S0406, and IRDye80 (39). In this work, we use OTL38, a folate analog conjugated to S0406 that targets the overexpression of cell surface folate receptors (FR) found in many types of cancers including ovarian, non-small-cell lung, mesotheliomas, and triple-negative breast cancers (40, 41).

In addition, noting that NIR fluorophores typically have lower fluorescence quantum yield than visible fluorophores, we re-designed our fiber probe assemblies to allow improved geometric light collection and rejection of instrument autofluorescence compared to our older design (29).

## 2 Materials and Methods

### 2.1 NIR-DiFC Instrument

The schematic of NIR-DiFC is shown in **Fig. 1a**. The light source is a tunable pulsed laser (Mai Tai XF-1, Spectra Physics, Santa Clara, CA) with excitation wavelength set to 770 nm. The power is adjusted with a variable attenuator (VA) before it is passed through a 766/13 nm bandpass clean up filter (BP-ex; FF01-766/13-25, IDEX Health and Science LLC, Rochester, NY). The light is then split into two beams with a beam splitter (BS; 49005, Edmund Optics, Barrington, NJ) before being coupled with a collimation package (FC-ex; F240SMA-780, Thorlabs Inc., Newton, NJ) into source fibers of the fiber probe assemblies (see Section 2.2 Integrated NIR-DiFC Fiber Probe Design). The light power at the sample is set to 25 mW. The output of the probe collection fibers is collimated (FC-em; F240SMA-780, Thorlabs) and the light is passed through an 810/10 nm bandpass emission filter (BP-em; FF01-810/10-25, IDEX Health and Science LLC) before being focused on to the surface of a photomultiplier tube (PMT; H10721-20, Hamamatsu, Bridgewater, New Jersey) with a 30 mm focal length lens (L-em; 67543, Edmund Optics). The PMTs are powered by a power supply (C10709; Hamamatsu). Output signals from the PMTs are filtered with an electronic 100 Hz low pass filter, amplified with a low-noise current pre-amplifier (PA; SR570, Stanford Research Systems, Sunnyvale, CA), and then acquired with a data acquisition board (USB-6343 BNC; National Instruments, Austin, TX). **Fig. 1b** shows a photograph of the instrument.

**Fig. 1.**
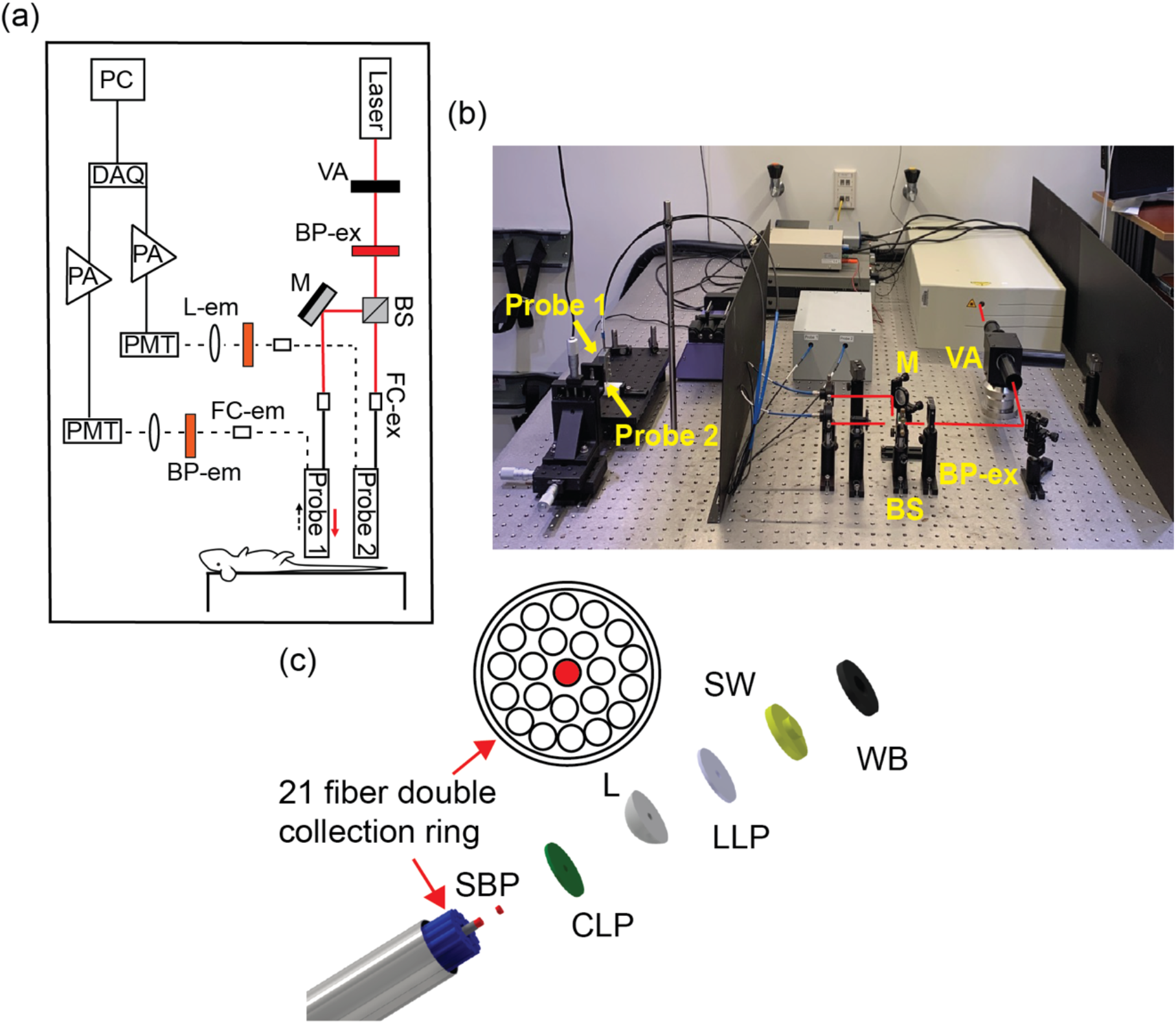
The NIR-DiFC instrument. (a) optical system schematic. (b) Photograph of the NIR-DiFC system with main components labeled (c) schematic of an NIR-DiFC fiber probe bundle. See text for component details.

### 2.2 Integrated NIR-DiFC Fiber Probe Design

NIR-DiFC uses custom-designed integrated fiber probe assemblies as shown in **Fig. 1c** (EmVision LLC, Loxahatchee, FL). The design is an improvement compared to our previous fiber probe design (29) with better geometric collection efficiency and autofluorescence suppression. Each probe is constructed with 21 all silica low hydroxyl (OH) content 300 µm core 0.22 NA collection fibers, arranged in a 7-fiber inner ring and 14 fiber outer ring. The 21 collection fibers are arranged around a single fiber for laser delivery which is also an all silica 300 µm core low OH, 0.22 NA fiber.

A donut-shaped 807 nm long-pass filter (CLP; BLP01-785R, IDEX Health and Science LLC) is positioned in front of the 21 collection fibers to reject laser light and pass collected fluorescence light from the sample. The source delivery fiber has a smaller 709/167 nm band pass filter (SBP; FF01-709/167, IDEX Health and Science LLC) positioned in front which allows the light to pass but rejects any other interfering light created from autofluorescence of the probe materials. The distal end of the probe also has a converging sapphire lens (L) with a hole drilled in the center, and a second lens long pass filter (LLP) with a hole in the center placed on the lens. The laser fiber passes through the center hole of the fiber filter, lens, and lens filter, which eliminates any reflected laser light from the lens surface which can create undesired autofluorescence from the probe materials. The LLP filter further reduces the laser light which can otherwise back reflect from the sample into the probe and create unwanted background fluorescence. A magnesium fluoride stepped window (SW) is attached to the lens, lens filter, and excitation fiber assembly. This stepped window allows the laser light and collection light paths to overlap at the distal window tip-sample interface, maximizing collection of fluorescent light. The stepped window and window block (WB) reject undesired light from being collected that is excited outside of the desired sample region and improves blocking efficiency of the long pass filters. The fibers, lens, and other optical components are placed inside a 2.4 mm outside diameter stainless steel needle tube.

### 2.2 Fluorescent Microspheres

We used “Jade Green Low Intensity” (JGLI; FL100782, Spherotech, Lake Forest, IL) NIR fluorescent microspheres. These microspheres are approximately 10-14 µm in diameter and (as we show) have similar fluorescence brightness to CTCs labeled with an NIR fluorescent molecular probe.

### 2.3 NIR-DiFC Testing in Tissue-Mimicking Flow Phantoms In Vitro

NIR-DiFC was first tested using a tissue-mimicking flow phantom (**Fig. 2a**) as we have used previously (18, 28, 30, 31, 36). The phantom is made from high-density polyethylene (HDPE) block and has similar absorption and scattering properties to biological tissue (18). Microbore Tygon tubing (TGY1010C, Small Parts, Inc., Seatle, WA) was inserted into a hole drilled into the phantom at a depth of 0.75 mm. Using a microsyringe pump (702209, Harvard Apparatus, Holliston, MA), 1 mL suspensions of microspheres or NIR fluorophore labeled cells (see Section 2.4 below) suspended in PBS at a concentration of 10^3^ per mL were flowed through the phantom at a flow rate of 50 µL/min.

**Fig. 2.**
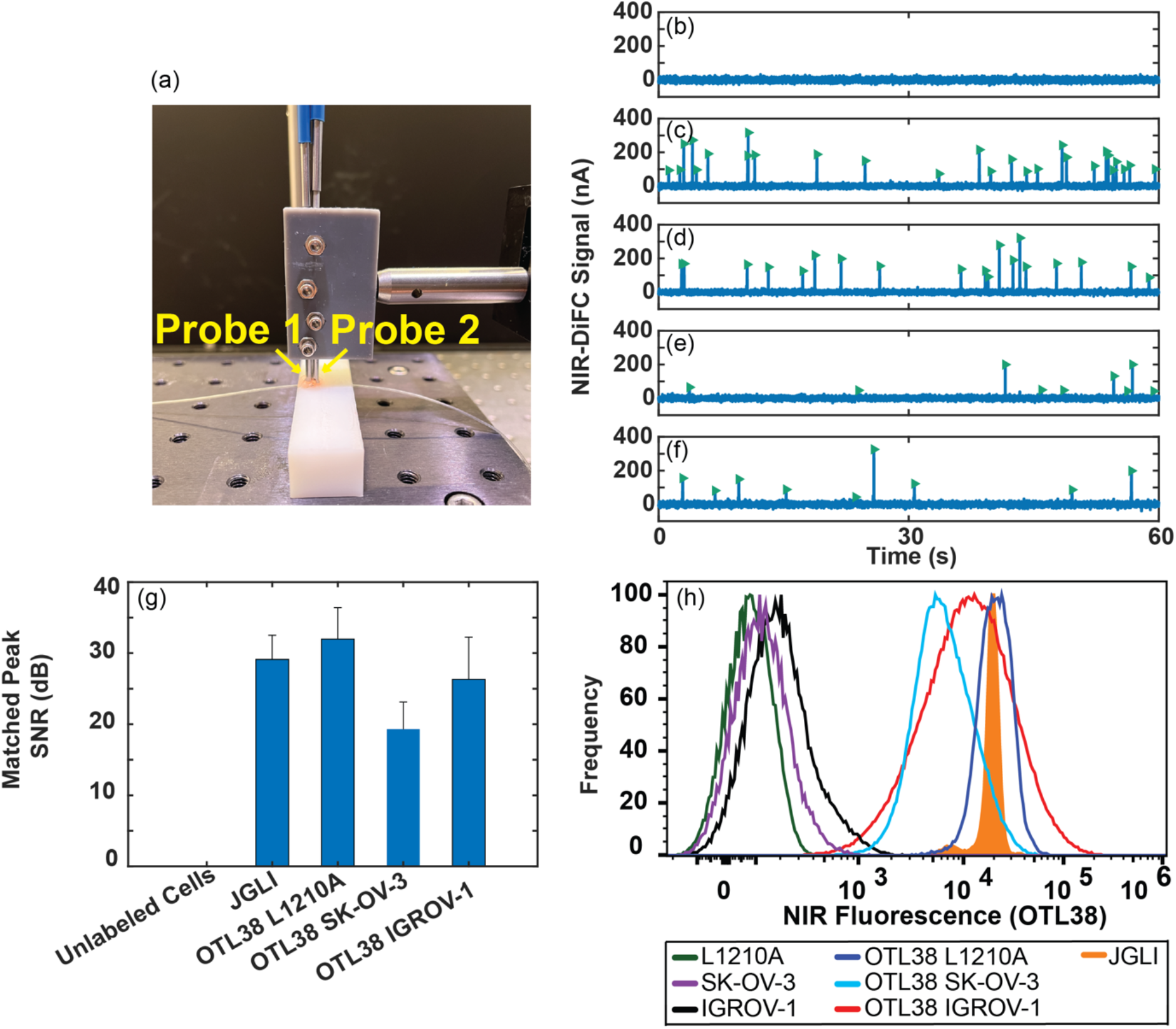
NIR-DiFC validation with (a) a tissue mimicking flow phantom model. The NIR-DiFC fiber probes are placed on the surface, approximately above a strand of Tygon tubing embedded 0.75 mm deep, through which we flowed suspensions of microspheres or cells. (b) after signal processing, no peaks were observed with control suspensions of unlabeled L1210A showing the stability of the system. Representative NIR-DiFC data collected from solutions of (c) JGLI microspheres, OTL38 labeled (d) L1210A cells, (e) SKOV-3 cells, and (f) IGROV-1 cells are shown. Each transient peak (arrowhead symbols) represents a forward-matched detection. (g) Comparison of the signal-to-noise (SNR) ratio of detected peaks for each sample. (h) Fluorescence intensity (measured by flow cytometry) of OTL38 labeled cells, unlabeled cells, and JGLI microspheres, showing good general agreement with NIR-DiFC fluorescence measurements.

### 2.4 NIR Fluorescent Labeling of Cells with OTL38

We used L1210A immortalized murine leukemia cells which were previously modified to over-express FR (42-44). We also used SK-OV-3 (Angio-Proteomie, Boston, MA) and IGROV-1 (Sigma Aldrich, St. Louis, MO) which are both naturally expressing FR+ immortalized human ovarian cancer cell lines. All cells were cultured in RPMI 1640 folic acid deficient media (Gibco 27016021; ThermoFisher Scientific, Waltham, MA) supplemented with 10% fetal bovine serum (Gibco 16000044; ThermoFisher Scientific), 1% penicillin/streptomycin (Gibco 15140122; ThermoFisher Scientific) and incubated at 37° with 5% CO_2_.

OTL38 (On Target Laboratories, West Lafayette, IN) is a FRα-targeting fluorescent small-molecule (MW 1414.42 Da) contrast agent that has previously been characterized in addition to recently being fully FDA approved for the use in fluorescence guided surgery (45-47). OTL38 is a conjugate between a folate analog and S0456 dye (similar fluorescence spectrum to ICG) with a maximum excitation wavelength of 776 nm and stokes shift of 17 nm (48).

1 mL suspensions of 10^6^ FR-over-expressing L1210A, SK-OV-3, or IGROV-1 cells in 2% FBS in PBS (Gibco 10010049; ThermoFisher Scientific) were co-incubated with 200 nM (20 µL of 10 µM stock) OTL38 at 37° with 5% C0_2_ for 1 hour. Cells were then washed twice with PBS before start of experiments.

### 2.5 Flow Cytometry

OTL38 was analyzed using a benchtop Attune NXT flow cytometer (ThermoFisher Scientific). NIR fluorescence was collected using a 637nm excitation wavelength and a 780/60nm emission filter.

### 2.6 Validation of NIR-DiFC in Mice In Vivo

All mice were handled in accordance with Northeastern University’s Institutional Animal Care and Use committee (IACUC) policies on animal care. Animal experiments were carried out under Northeastern University IACUC protocol #21-0412R. All experiments and methods were performed with approval from and in accordance with relevant guidelines and regulations of northeastern University IACUC.

For proof-of-concept testing of NIR-DiFC *in vivo*, we injected L1210A cells intravenously in nude mice. We and others have previously shown that these circulate in stable numbers in the blood for hours after injection (21, 36). L1210A cells were labeled with OTL38 *in vitro* as described above. A suspension of 10^6^ labeled cells in 100 µL cell culture media was injected *i.v*. via the tail vein (N = 3) of 6–8-week-old female Athymic nude mice (Athymic NCR Nu/Nu/553; Jackson Laboratory, Bar Harbor, ME). NIR-DiFC was preformed 10 minutes after injection on the ventral caudal tail artery for 60 minutes.

### 2.7 DiFC Data Analysis

We used the DiFC signal processing approach reported in detail by us previously (28). Briefly, NIR-DiFC data was acquired, and the moving mean background signal subtracted from the data. This background signal results from autofluorescence of the biological tissue as well as intrinsic autofluorescence of the instrument, for example from optical fibers, filters etc. The noise on this background (post mean-subtraction) is inherent to the DiFC measurement. This necessitates the use of a detection thresholds, which we define as 5 times the standard deviation of the background signal (calculated dynamically for each minute of the scan). Measured transient fluorescent peaks from above this threshold are considered “peak candidates”. To further discriminate system noise, we impose the constraint that two peaks must be detected sequentially between the first and second probes with a time delay consistent with the separation distance of the two probes and the estimated speed of the cell, which is calculated from the peak width. We term these peaks “forward matches” and correspond to cells traveling (in this case) in the ventral caudal arteries. Other unmatched (capillary), reverse-matched (venous) or spurious peaks due to noise were rejected from the analysis.

## 3 Results

### 3.1 NIR-DiFC Detection of Microspheres and Cells in Flow Phantom In Vitro

We first tested NIR-DiFC using an optical flow phantom (**Fig. 2a**), which we have shown previously approximates the optical properties of bulk biological tissue in the NIR range (35). Peaks were never observed in control solutions of either PBS or unlabeled cells; a representative NIR-DiFC data scan for unlabeled L1210A cells is shown in **fig 2b**. We passed suspensions of fluorescent microspheres and OTL38 labeled cells through the phantom to mimic cells flowing through a blood vessel. Representative NIR-DiFC data (background subtracted) is shown in **Fig. 2c-f** for JGLI microspheres, L1210A cells, SKOV-3 cells, and IGROV-1 cells, respectively. In each plot, every vertical peak represents a forward-matched (between the two NIR-DIFC probes) cell or microspheres. Differing peak amplitudes are due primarily to the variations in OTL38 contrast agent uptake by the cells. The standard deviation of system noise in phantoms was α = 7.9 +/- 0.1 nA.

The signal to noise ratio (SNR) for the different microsphere and cell types, averaged over a minimum of 90 peaks is summarized in **Fig. 2g**. Here, SNR is defined as 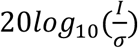, where I is the mean peak amplitude. The relative detection intensities are approximately in agreement with those measured with flow cytometry as shown in **Fig. 2h**.

### 3.2 In Vivo validation of NIR-DiFC with OTL38 labeled L1210A cells

We next tested NIR-DiFC in nude mice *in vivo*. FR+ L1210A cells were pre-labeled with OTL38 *in vitro* and then injected *i.v*. via the tail vein. The fiber probes were placed on the skin surface on the ventral side of the tail, approximately above the ventral caudal vascular bundle as shown in **Fig. 3a**. Representative data from an OTL38-L1210A injected mouse are shown in **Fig. 3b**. These peaks were never observed in control (un-injected) mice as shown in **Fig. 3c**.

**Fig. 3.**
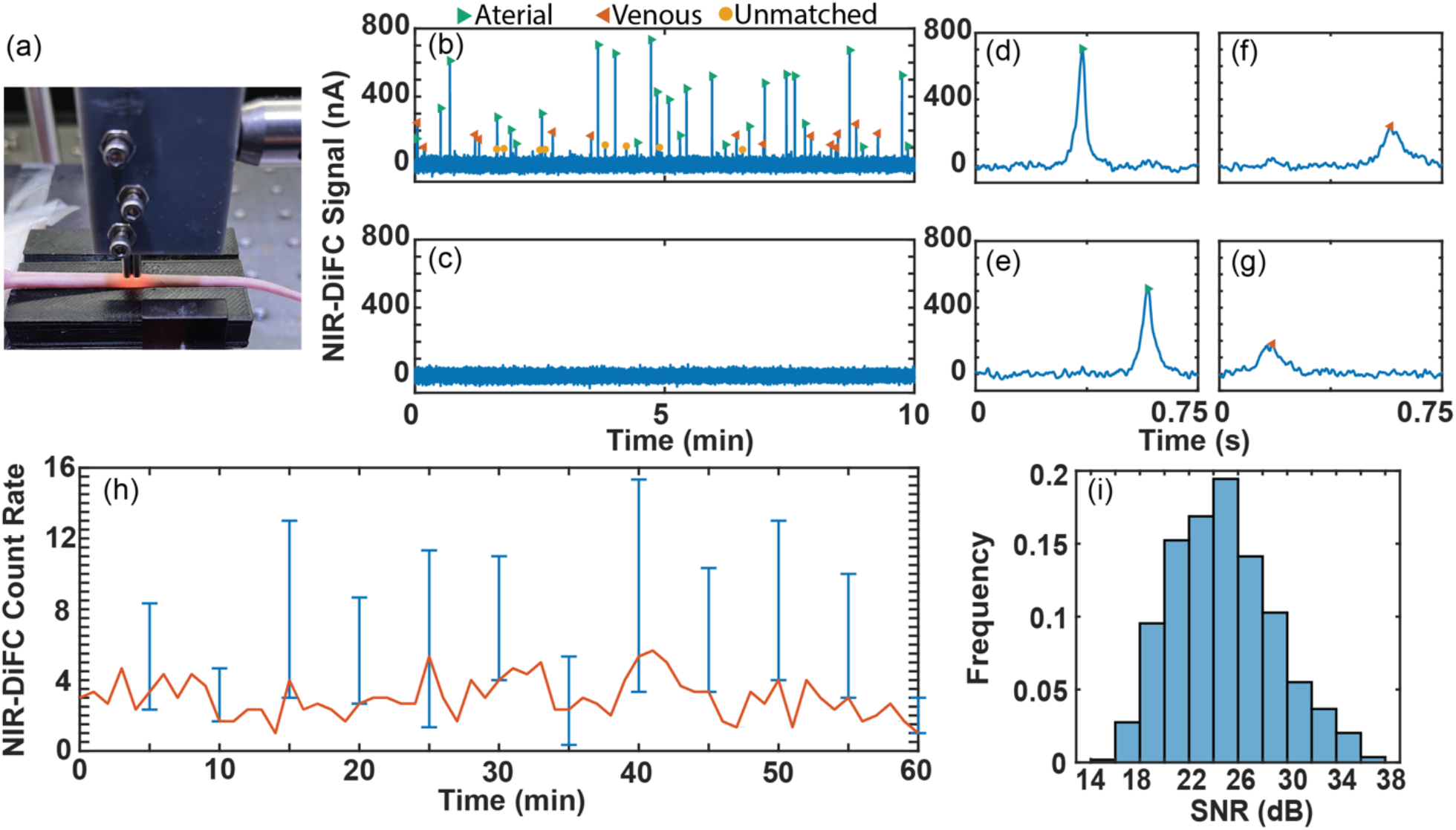
*In vivo* validation of NIR-DiFC. (a) Fiber probes were placed on the surface of the mouse tail approximately above the ventral caudal artery. (b) Representative NIR-DiFC data measured from a mouse injected *i.v*. with OTL38 labeled L1210A cells. Each green arrowhead is an arterial matched cell (traveling in artery), each orange arrowhead is a venous matched cell (traveling in vein) and each yellow labeled peak represents a detected unmatched cell (traveling in capillary). (c) Representative NIR-DiFC data from an un-injected control mouse. (d,e) Cell detected sequentially in probes 1 and 2, indicating a cell traveling in the arterial direction. (f,g) Cell detected sequentially in probes 2 and 1, indicating a cell traveling in the venous direction. (h) The mean cell detection rate over a 1-hour scan indicating that cell numbers were approximately stable in circulation. Range bars shows the minimum and maximum values (N = 3). (i) Histogram of detected peak SNRs.

As described in Section 2.7 above, for *in vivo* measurements we imposed a matching condition, examples of which are shown in **Figs. 3d-g**. Cells traveling in the forward, arterial direction are detected as peaks sequentially measured on the first (**Fig. 3d**) and then second (**Fig. 3e**) probes and separated by a time delay due to the 3 mm separation between probes. These are indicated as green forward arrow markers in longer data trace in **Fig. 3b**. Likewise, cells traveling in the reverse (venous) direction are sequentially detected by the second and then first probe (**Figs. 3f-g**) are indicated as orange reverse arrows. As shown and as we noted previously cells moving in the arterial direction (forward) are generally faster moving (shorter period between sequential detections) than those moving in the venous direction (reverse) (28). Unmatched peaks - likely due to cells traveling in the network of capillaries – are indicated as round markers.

As shown in **Fig. 3h**, the cell detection count rate stayed relatively constant over 60-minute scans, which is consistent with our previously observed circulation kinetics in the same mouse model and cell line (36). A histogram of signal-to-noise ratios (SNR) of all detected peaks over all mice measured in this study is shown in **Fig. 3i**. The mean peak SNR was 26 dB. We previously used a green folate targeted molecular probe (EC17) with the same mouse model and found a mean peak SNR of 18 dB. While we note there are differences in the instrument, fluorophores, placement of the probe (tail versus hindleg), it is noteworthy that the SNR was significantly improved compared to our blue-green (b-DiFC) system.

### 3.3 NIR-DiFC Autofluorescence is Significantly Lower than Visible DiFC

In addition to the advantages of NIR-DiFC described above, as summarized in **fig. 4** we observed that tissue autofluorescence is markedly reduced and the signal is free of motion artifacts (35) compared to our pre-clinical GFP-compatible b-DiFC instrument (31). Use of blue laser light is widely understood to yield higher biological tissue autofluorescence than near-infrared light (49). Likewise, use of blue light in fluorescence measurements is in general more sensitive to breathing (motion) or photoplethysmography artifacts (50-52). We occasionally observed this in our prior work with b-DiFC: an example b-DiFC data trace showing a breathing artifact is shown in **fig 4a** (18, 30). Generally, when breathing artifacts are observed we simply adjust or re-position the b-DiFC fiber probe on the mouse skin surface to remove it (**fig. 4b)**. However, these artifacts may occasionally result in false-positive peak detections.

**Fig. 4.**
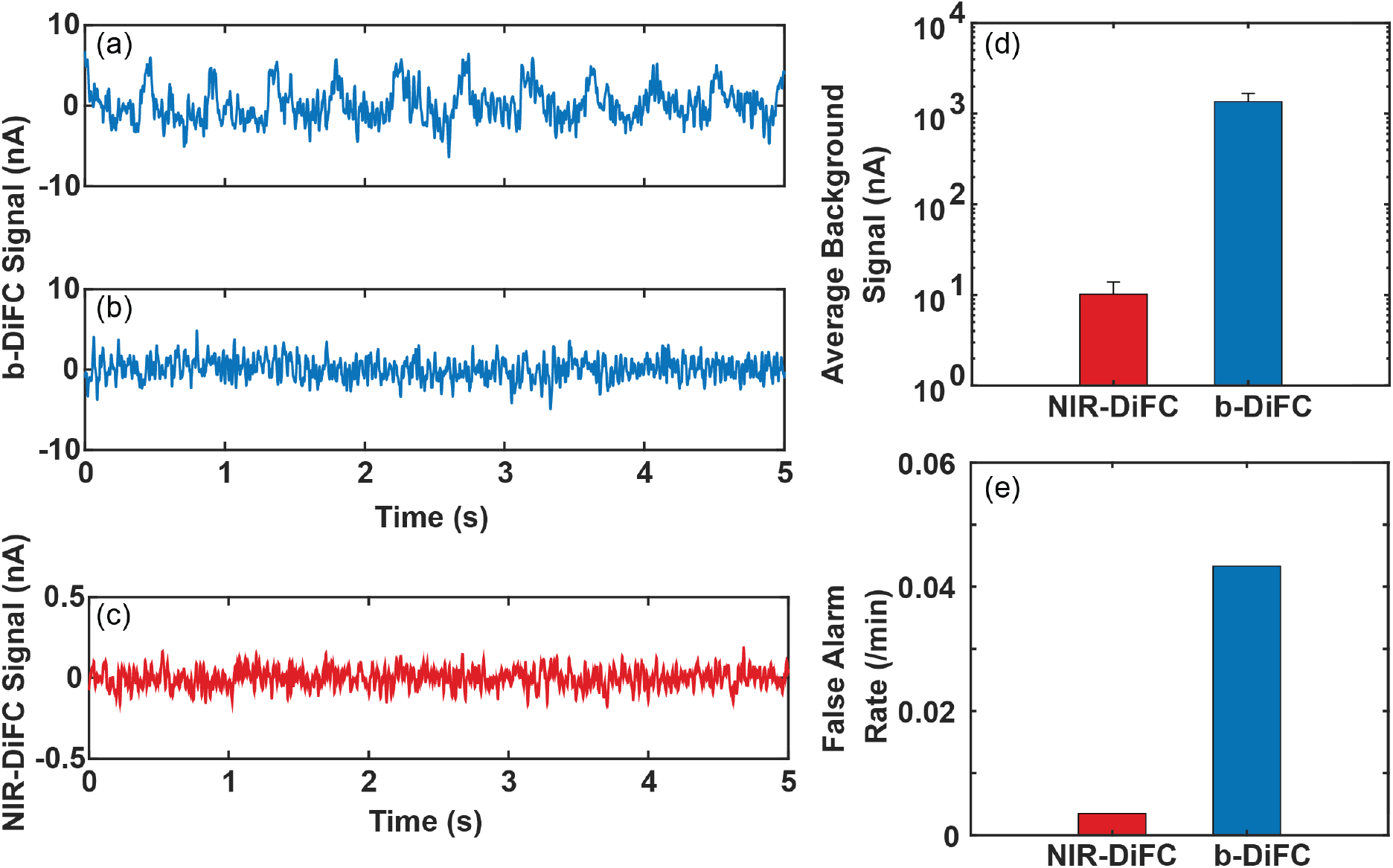
(a) Example breathing artifacts measured with our blue-green b-DiFC instrument. (b) These generally can be corrected by simply adjusting the b-DiFC probe position on the skin, although this causes loss of time. (c) In contrast, these motion (breathing) artifacts were not observed with NIR-DiFC measurements. (d) Use of NIR laser light for DiFC resulted in significantly lower tissue autofluorescence than blue light. (e) In practice, this resulted in a reduction in DiFC false alarm rate.

In contrast, these motion artifacts were not observed in NIR-DiFC (e.g. **fig. 4c**) which we attribute to lower sensitivity to motion artifacts and generally reduced levels of tissue autofluorescence (33). This is illustrated in **fig. 4d**, which shows the mean tissue autofluorescence for b-DiFC and NIR-DiFC systems. The laser power at the surface was 20 and 25 mW for the b-DiFC and NIR-DiFC respectively. We note that in making this comparison we explicitly standardized (using the Hamamatsu H10722-20 PMT specifications documentation) the output current between the two systems correcting for voltage output (b-DiFC) and current output (NIR-DiFC) versions of the PMT module and correcting for differences in control voltage applied between PMTs (which were lower with the b-DiFC system than NIR-DiFC system). In practice, as shown in **Fig. 4e**, lower noise levels resulted in a lower false alarm rate (FAR) with the NIR-DiFC system compared to the b-DiFC system.

## 4 Discussion and Conclusion

We have previously reported red and blue-green versions of DiFC, which we used to enumerate fluorescently labeled or fluorescent protein expressing CTCs during metastasis development in several mouse models (18, 30, 31). Beyond small animal models, one long-term goal for DiFC is potential use in humans (32). As a first step to this, here we reported development of a NIR version of DiFC. The rationale was three-fold. First, NIR-DiFC should allow detection of CTCs in blood vessels 2-4 mm deep using our DiFC approach (35). Hence, major blood vessels in the wrist or forearm of a human should be accessible with NIR-DiFC (34).

Second, use of NIR laser light for DiFC yields lower tissue autofluorescence and lower incidence of motion artifacts in DiFC data (**fig. 4**). In practice this results in fewer false positive CTC detections and obviates the need for occasional re-aligning of the instrument. NIR fluorophores are typically less bright than visible fluorophores, but as we showed the improved NIR-DiFC fiber probe design allowed efficient geometric collection of fluorescent light. Combined with lower autofluorescence, we showed clear detection of CTCs in vivo with SNRs approximately 8 dB higher than analogous experiments with our b-DiFC system (36).

Third, many emerging fluorescence molecular contrast agents use NIR fluorophores because they allow imaging in deeper tissues clinically (39).As such, we designed NIR-DiFC to be compatible with these. We are currently evaluating OTL38 for labeling of CTCs for NIR-DiFC. We and others previously showed that small-molecule folate receptor targeted probes have significant potential for bright and specific labeling of CTCs in whole blood (36). This is an ongoing area of research in our group.

Future work will evaluate labeling and detection of CTCs by systemic administration of OTL38 *in vivo* (as opposed to pre-labeling *in vitro* as we have done here) in pre-clinical metastasis models. Alternative NIR contrast agents may be tested in the future, as well as use of NIR-DiFC for other types of rare circulating cells.

## Acknowledgements

This work was funded by the National Institutes of Health (R21CA246413). The authors thank Prof. Philip S. Low and Dr. Madduri Srinivasarao of Purdue University, West Lafayette, IN, for the generous gift of OTL38 contrast agent used in this work.

## Code, Data, and Materials Availability

The datasets generated and analyzed during this work and the MATLAB code are available through the Pennsieve data sharing platform DOI: 10.26275/h0fa-hd8z. Additionally, MATLAB processing code can be found in a Niedre Lab Github repository: https://github.com/mark-niedre/Design-and-Validation-of-NIR-DiFC (DOI:10.5281/zenodo.6506804).

**Joshua Pace** received his BS degree in biomedical engineering from the University of Arizona in 2019. He is currently a PhD Candidate in the Northeastern University Department of Bioengineering.

**Mark Niedre** received his PhD from the University of Toronto Department of Medical Physics in 2004. He is a Professor of Bioengineering at Northeastern University and a senior member of SPIE.

